# SPLASH2 provides ultra-efficient, scalable, and unsupervised discovery on raw sequencing reads

**DOI:** 10.1101/2023.03.17.533189

**Authors:** Marek Kokot, Roozbeh Dehghannasiri, Tavor Baharav, Julia Salzman, Sebastian Deorowicz

**Author notes:** These authors contributed equally to this work (Marek Kokot, Roozbeh Dehghannasiri). Eric and Wendy Schmidt Center, Broad Institute, Cambridge, MA 02142, USA, Department of Data Science, Dana-Farber Cancer Institute, Boston, MA 02115, USA.

## Abstract

SPLASH is an unsupervised, reference-free, and unifying algorithm that discovers regulated sequence variation through statistical analysis of *k*-mer composition, subsuming many application-specific methods. Here, we introduce SPLASH2, a fast, scalable implementation of SPLASH based on an efficient *k*-mer counting approach. SPLASH2 enables rapid analysis of massive datasets from a wide range of sequencing technologies and biological contexts, delivering unparalleled scale and speed. The SPLASH2 algorithm unveils new biology (without tuning) in single-cell RNA-sequencing data from human muscle cells, as well as bulk RNA-seq from the entire Cancer Cell Line Encyclopedia (CCLE), including substantial unannotated alternative splicing in cancer transcriptome. The same untuned SPLASH2 algorithm recovers the BCR-ABL gene fusion, and detects circRNA sensitively and specifically, underscoring SPLASH2’s unmatched precision and scalability across diverse RNA-seq detection tasks.

## Text

Regulated RNA transcript diversification can occur through myriad mechanisms that include alternative splicing, RNA editing, and the inclusion of sequences absent from the reference genome. Massive RNA-seq datasets provide an unprecedented opportunity to discover regulated transcriptomic complexity in healthy tissues as well as its implications for disease. Despite the promise of computational approaches to identify targetable transcript dysregulation, methods to analyze these datasets face limitations, including computational time constraints and algorithmic biases such as in read alignment. Alignment, while useful for many applications, could lead to missing entire families of splicing events such as circRNAs (Salzman et al. 2012) and could censor reads representing important mechanisms, particularly in biological contexts such as tumors that a reference genome cannot sufficiently model. Moreover, alignment is computationally demanding and poses storage issues, with alignment output files often exceeding input raw reads by a factor of two, hindering the processing of massive datasets. Moreover, alignment serves as only the initial step in workflows seeking to infer differential regulation of RNA variation. After alignment, users must navigate through a plethora of time-consuming bioinformatic algorithms to detect each specific signal such as RNA splicing, RNA editing, and V(D)J recombination, diversifying immune repertoires.

We recently introduced SPLASH (Chaung et al. 2023) which leverages a unified statistical framework to detect sample-regulated RNA sequence variation by analyzing the *k*-mer composition of reads. At its core, SPLASH finds constant sequences (anchors) that are followed by a set of diverse sequences (targets) with sample-specific expression distribution by providing valid p-values (Chaung et al. 2023). SPLASH is reference-free, sidestepping the computational challenges associated with alignment and making SPLASH in principle faster than other alignment-based methods, enabling discoveries that would have been otherwise impossible. However, the original implementation of SPLASH still faced limitations in processing large datasets, such as the one we highlight in this study, 550GB of raw reads from a study of Amyotrophic Lateral Sclerosis disease (Ma et al. 2022).

Here, we introduce SPLASH2 (https://github.com/refresh-bio/SPLASH) an ultra-fast, memory-efficient, and user-friendly implementation of SPLASH. SPLASH’s initial step involves counting *k*-mer tuples of anchors and targets, a process that was highly time-consuming and memory-intensive in the original implementation, making it impractical for large datasets. SPLASH2 resolves this by leveraging KMC (Deorowicz, Debudaj-Grabysz, and Grabowski 2013; Kokot, Dlugosz, and Deorowicz 2017), a highly efficient *k*-mer counter well-suited for processing large-scale genomic datasets through memory optimization and architecture parallelization techniques.

SPLASH2 is implemented in 3 main steps (Figure 1A). In step 1, SPLASH2 parses input FASTQ files and counts anchor-target pairs in each sample separately. If gaps are desired between an anchor and its target sequences, they are subsequently removed during parsing. The output of step 1 is a list of lexicographically sorted records containing anchor, target, and count for each sample. Step 2 involves merging the target sequences and their counts found for each anchor across all samples to build a contingency table of target counts for each unique anchor. The contingency table is then used to compute a statistically valid p-value through an unsupervised optimization (Baharav, Tse, and Salzman 2024; Chaung et al. 2023) (Methods). This step is memory-frugal as the contingency table is represented by a sparse matrix and each contingency table is loaded into memory individually. Finally, multiple testing correction is performed in Step 3. To enhance the interpretability of SPLASH2’s calls and classify each anchor into distinct biologically relevant categories, we construct *extendor* sequences by concatenating anchor-target pairs. Similar to the classification approach for SPLASH (Dehghannasiri et al. 2022), we then align extendors to the reference genome and classify each anchor using the alignment information of the two extendors corresponding to the top two most abundant targets for the anchor.

**Figure 1.**
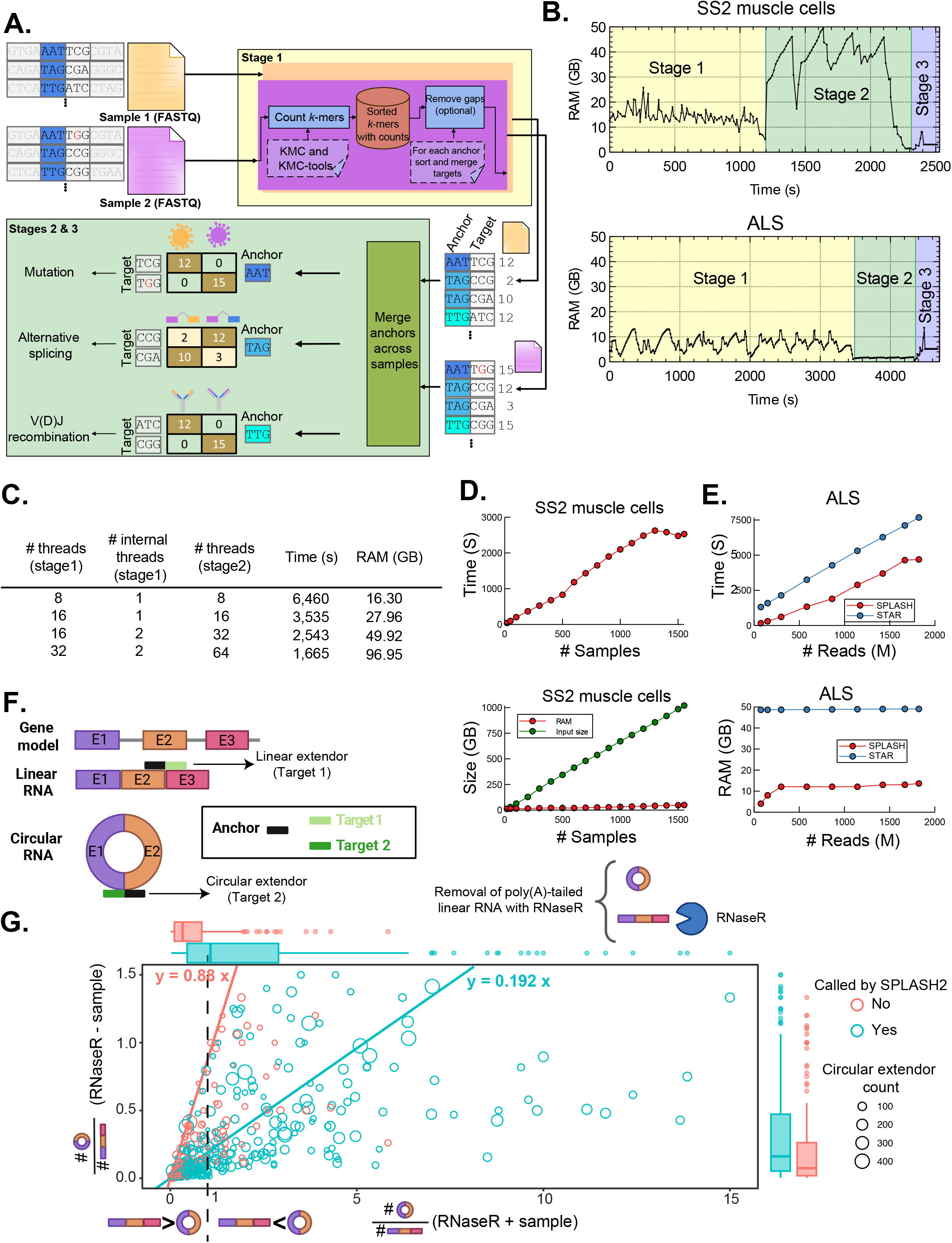
SPLASH2 pipeline outline and evaluation. (A) Outline of the SPLASH2 pipeline involving parsing FASTQ files to extract anchor-target pairs and their counts (Stage 1), merging targets and their counts across samples and computing p-values (Stage 2), and multiple testing correction (Stage 3). Anchors with significant p-values are classified into biologically relevant events such as alternative splicing. (B) Memory and run-time requirements are shown for SS2 muscle cells (top plot) and the ALS dataset (bottom plot), indicating the impact of sample size and sequencing depth on the computational burden of Stages 1 and 2. (C) Time and memory requirements for analyzing SS2 muscle cells with varying numbers of threads illustrate the trade-off between computation speed and RAM usage. (D) Time (top) and memory (bottom) requirements increase linearly with the number of input samples for SS2 muscle cells. (E) Run time (top) increases linearly with input reads, while memory usage (bottom) saturates at 15GB for the ALS dataset. SPLASH2 demonstrates at least 1.7 times faster and 4 times more memory-efficient performance compared to STAR. (F) Detection of circular RNAs through anchors with one extendor mapping to the back-splice junction and one extendor mapping to the canonical junction. (G) Scatter plot showing the disparity in count ratios between circular and linear extendors in RNase R +/- samples of each cell line for SPLASH2-called circRNA anchors (green) versus uncalled anchors (red). Circle size represents circular counts and linear regressions were weighted by circular extendor count. Each box shows 25-75% quantiles of circular-to-linear count ratios in RNase R+ (horizontal), RNase R - (vertical), and for anchors called (cyan) and not called by SPLASH2 (red). The midline of each box is the median and whiskers extend to 1.5 times the interquartile range. All points outside of this range are plotted individually.

We characterized the time and memory requirements for each step of SPLASH2 using 1,553 human muscle single cells (3.7 billion reads) profiled with SmartSeq2 (SS2) (Tabula Sapiens Consortium* et al. 2022) and an Amyotrophic Lateral Sclerosis (ALS) dataset (1.8 billion reads; (Ma et al. 2022)) (Figure 1B). Step 2, involving merging target counts across samples, required more time and memory when analyzing SS2 (many samples with fewer reads per sample), while Step 1, involving parsing each sample for extracting anchor and target sequences, required more resources for the ALS dataset (few samples but deeper sequencing). Step 3 accounted for < 10% of the total run time (Figure 1B). As our implementation supports multithreading, it offers reduced run time but at the cost of increased memory consumption (Figure 1C). Memory usage and runtime grow linearly with input size (Figure 1D), except for RAM usage for ALS (Figure 1E). SPLASH2 was at least 1.7 times faster and consumed 4 times less memory (Figure 1E) compared to STAR (Dobin et al. 2013). This is particularly noteworthy considering that STAR alignment is only the initial, relatively less intensive step in typical custom bioinformatics analyses. SPLASH2 detects cryptic splicing in positive controls *UNC13A, STMN2*, and *POLDIP3* experimentally validated for functional roles in ALS; beyond these examples, SPLASH2 predicted novel cryptic splicing events missed by other methods (Ma et al. 2022). Our analysis indicated minimal impact on run time, RAM usage, and significant anchors when varying SPLASH’s parameters (i.e., anchor, target, and gap length and fraction of training data), except for moderate values of anchor length between 10 and 20 (Suppl. Figure 1).

We benchmarked SPLASH2 using data from a study of circular RNAs (circRNAs) (Vromman et al. 2023). This dataset contains 3 cell lines (colon, lung, hepatocellular), each with two samples: one treated with RNase R+, which enriches for circular RNAs by enzymatic digestion of linear RNA, and one untreated (RNase R-). We ran SPLASH2 separately on the RNase R+/- samples for each cell line (Methods). We define a circRNA anchor if one extendor maps to a back-splice junction (or a junction between exons in non-increasing order), a unique splicing pattern defining circRNAs, and the other extendor maps to the linear RNA in the same locus (Figure 1F). SPLASH2 deems a circRNA anchor significant (and consequently we define that it identifies the corresponding circRNA) if it finds sufficient statistical evidence for the differential representation of circ- and linear RNA targets between RNAse R+ and RNase R- within a cell line. K-mer preprocessing preceding SPLASH2 identified anchors for 1,265 out of 1,560 validated circular RNAs across three cell lines, surpassing other circRNA-specific tools (Suppl. Figure 2A, Suppl. Table 1). However, SPLASH2 calls an anchor– in this case, circRNA anchor– when it is present in the sample, and also its target (corresponding to the circRNA) is differentially regulated between samples compared to targets corresponding to other alternative splice variants. Of 1,265 initially detected circRNAs, 521 were called significant by SPLASH2 (Suppl. Figure 2 A, Suppl. Table 1, Methods) that have the added quality of being differentially represented as compared to their linear counterparts in RNase R treatment. Notably, significant circRNA anchors showed a significantly larger difference in circular-to-linear ratios between RNase R +/- samples compared to non-significant anchors (Figure 1G). Together, these results suggest SPLASH2’s robust performance in circRNA detection, despite not being tailored or “prompted” for circRNA detection. SPLASH2 not only exhibits sensitivity comparable to other tools but also uniquely identifies those sequences with differential representation between conditions that enrich for circRNA (RNAse R +) and those that do not (RNAse R -), a quality missing in other benchmarked tools (Vromman et al. 2023).

We also evaluated SPLASH2’s specificity using the circRNA benchmarking study’s amplicon sequencing data, which amplified a selected panel of 1560 circRNAs (validated circRNAs) with PCR followed by sequencing from the same cell lines. This data lacks true ground truth and potentially contains additional circRNAs due to predicted off-target amplification (Vromman et al. 2023) or resulting in more than one circRNA junction due to alternatively spliced variants in amplicon data (Salzman et al. 2013; Chen et al. 2023), but offers a relatively reliable basis for specificity analysis. SPLASH2 detected 1,852 circRNAs; 92% were either validated circRNAs or had at least one annotated exon boundary (Suppl. Figure 2B, Methods), suggesting a biological origin for these differentially spliced circRNAs, a known mechanism of back-splicing (Salzman et al. 2013; Chen et al. 2023). For the remaining 148 circRNAs, we further assessed their proximity to detected circRNAs in original cell lines. 71 were close to a detected circRNA by other circRNA tools in the corresponding cell lines, defined as a distance of <20 base pairs. Among the other 77 circRNAs, only 3 circRNAs had a distance of at least 100bps to the closest annotated exon boundary or detected circRNA. However, when blatted, their sequences all had perfect alignment to the reported back splice junctions (Suppl. Figure 3). The majority of the aforementioned 148 circRNAs, 122 (82.4%) and 125 (84.5%), had detected circRNAs closer than the nearest annotated exon boundary based on 5’ and 3’ splice sites, respectively (Suppl. Figure 2C). Together, these findings show SPLASH2’s precision in detecting circRNAs without tuning or “task-specific” instructions, as required by other tools, and illustrate the general discovery power of SPLASH2 even within the realm of splicing, capturing linear and circRNA splicing detection in one algorithm.

We now demonstrate new biology unveiled by SPLASH2’s computational improvement. Our previous analysis of single-cell data was limited by computational constraints in SPLASH (Dehghannasiri et al. 2022), allowing analysis of only up to 400 cells with subsampled reads. SPLASH2 overcomes this hurdle, enabling simultaneous analysis of all 1,553 muscle cells from donor 2 (TSP2) in the Tabula Sapiens dataset (Tabula Sapiens Consortium* et al. 2022).

First, we show that SPLASH2, “unprompted” and untuned, rapidly identifies candidate RNA editing events *de novo*, including detecting potentially hyper-edited events, a task existing bioinformatic tools cannot perform efficiently (Figure 2 A, B). We selected anchors with at least four uniquely mapping extendors to the human reference genome, each comprising >5% of anchor reads, where the two most abundant targets exhibit single-base mismatches (Methods). The probability of such events occurring under a sequencing error model is vanishingly small, given that each cell in the dataset is sampled from a donor, implying a common genetic background. SPLASH2’s analysis revealed 402 anchors from 303 genes, where the targets of each anchor differ by single-base changes and fall into specific mismatch categories (Figure 2C, Suppl. Table 2). For example, the A<->G mismatch category means that all mismatches between targets can be resolved by converting A to G. The A<->G (or equivalently its reverse complement T<->C) mismatch category had ∼10-fold more anchors than other mismatch categories (A<->T, A<->C, G<->C), consistent with canonical A-to-I editing (Nishikura 2016). A<->T is the next most prevalent category, potentially explained by other known mechanisms (Uzonyi et al. 2021). Among the anchors categorized as A<->G mismatch with >3 mismatch positions in their dominant targets, those mapped to *AGO2, TDRP*, and *PECAM1 were* detected in 32 (2.1%), 15 (1%), and 12 (1%) cells, respectively, were found in the most cells (Figure 2D). The detected editing in *AGO2* supports our previous reports of editing (Dehghannasiri et al. 2022). We identified five mismatch positions, including one annotated SNP (according to dbSNP155) in the sequence of an anchor classified as G<->A in the 3’ UTR of *PECAM1*, a regulator of endothelial junctional integrity (Privratsky and Newman 2014). In three positions, targets diverge consistently with A-to-I editing but we cannot speculate about a causative mechanism for the other positions (Figure 2D).

**Figure 2.**
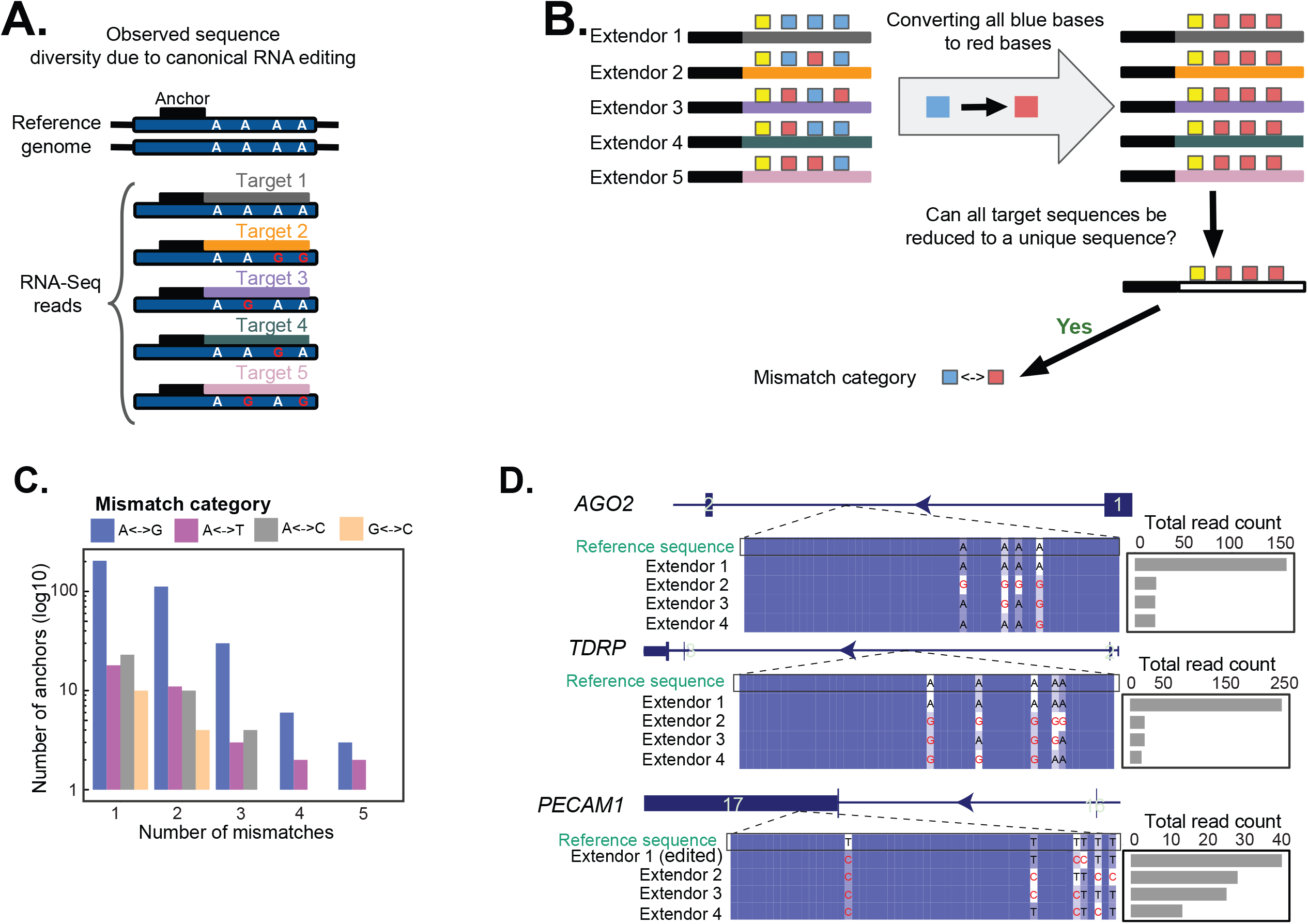
Rapid *de novo* discovery of RNA editing in scRNA-Seq by SPLASH2. (A) Diagram illustrating canonical RNA editing, where specific A’s in the reference sequence are converted to G’s in RNA transcripts, generating sequence diversity. (B) Toy example showing our approach for identifying anchors corresponding to specific RNA mismatch categories. (C) Number of anchors detected by SPLASH2 in SS2 muscle cells for each mismatch category and number of mismatches (in total: 512 anchors from 303 genes). The G<->A category has the highest number of anchors, consistent with canonical RNA editing. (D) Among genes with >3 mismatches, *AGO2, TDRP*, and *PECAM1* were identified in the highest number of cells. For each gene, multiple sequence alignment for the reference and the four most abundant extendors is provided, along with bar plots showing total read counts for each extendor across expressing cells. Gene models are based on CAT/liftoff annotation in T2T human genome reference assembly, with boxes representing exons (exon numbers indicating their sequential order in the gene) and lines introns and the number inside each exon shows its order among all exons in the gene.

SPLASH2 also enables the detection of both annotated and novel alternative splicing. An anchor is classified as unannotated alternative splicing if at least one STAR-mapped junction in the two most abundant extendors was not annotated in the human reference genome (T2T CAT/Liftoff annotations, Methods). In SS2 muscle cells, SPLASH2 identified 13,786 annotated and 11,584 unannotated cell-regulated (i.e., anchors with significant single-cell-dependent target distribution) splicing anchors (Suppl. Table 3). Among unannotated splicing anchors, anchors mapping to *SNHG17*, a noncoding RNA with reported roles in tumorigenesis (Pan et al. 2020), *CAMLG*, a calcium-modulating cyclophilin ligand, and *DIMT1*, an RNA-modifying enzyme with a role in ribosome biogenesis (Pan et al. 2020; Shen et al. 2021) were found in the highest number of cells (120, 118, and 112 cells, respectively) (Suppl. Figure 4).

To further highlight SPLASH2’s discovery power, we analyzed unannotated alternative splicing in the entire 671 primary cancer cell lines (5.7 TB) from primary tumors in the Cancer Cell Line Encyclopedia (CCLE), a comprehensive transcriptomic dataset, widely used for functional cancer biology (Ghandi et al. 2019). SPLASH2 took ∼47 hours with 50 Gb memory to process the entire corpus, while the STAR alignment alone (absent any custom downstream statistical or bioinformatic processing) took ∼80 hours (with the same memory) and required ∼4 TB of storage for the resulting BAM files. SPLASH2 reduced the number of input reads for STAR alignment from 64 billion ∼ 100-bp reads in CCLE FASTQ files to 30 million 54-bp extendors (2000-fold reduction). We also compared the efficiency improvement of SPLASH2 relative to the original implementation of SPLASH (Chaung et al. 2023): while the original SPLASH needed 26 hours to process just two CCLE samples, SPLASH2 finished the same task in 3.5 minutes (Methods). Previous analyses of CCLE have entailed separate bioinformatic approaches to profile mutations, splicing, or other events. Apart from being fragmented, these alignment-first analyses are expected to suffer from poor sensitivity to unannotated splicing, especially in cases involving mutations in exonic sequences, high mutation rates, or mapping to repetitive regions – known sources of mapping failure (Castel et al. 2015). These events are crucial to tumorigenesis and are likely to provide potentially valuable biomarkers and neoantigens (Stanley and Abdel-Wahab 2022; Quesada et al. 2011; Sveen et al. 2016).

Growing evidence suggests that many tumors acquire mutations in the spliceosome, leading to the activation of cryptic alternative splicing programs that contribute to pathology (Liu and Rabadan 2021). Thus, accurate detection of cryptic splicing in tumors holds paramount importance. This problem is particularly challenging due to the mutational burden and the accumulation of structural variants absent from the reference genome, some of which can be extensive (Korbel and Campbell 2013). 11,226 distinct alternative splicing events in 6,014 genes (excluding intron retention) were detected by SPLASH2 across the entire CCLE dataset. Establishing alternative splicing annotation categories using reported splice junction for each splicing anchor (Suppl. Figure 5 A) reveals that while the majority of the alternative splicing events (59% of alternative splicing events and 85.6% of their associated reads) correspond to annotated alternative splicing (Suppl. Figure 5 B, C, D), SPLASH2 detected extensive unannotated alternative splicing events in cell lines (Figure 3A), where 62.8% of the unannotated AS events involve one splice junction with annotated exon boundaries (the most prevalent unannotated AS category). 2,356 genes had at least one unannotated alternative splicing event (with >10% of anchor reads). Among the genes with unannotated alternative splicing, 199 were cataloged in the COSMIC database (Tate et al. 2019), a database of genes with known roles in tumorigenesis, implying an enrichment of COSMIC genes in SPLASH2 calls (exact binomial p-value = 1.92e-6, Methods). Notably, COSMIC genes *CTNNA2, NSD2, PTEN*, and *COL1A1* had the highest effect size. *PTEN* is a classic tumor suppressor (Cristofano, Di Cristofano, and Pandolfi 2000), and its loss has been linked to resistance to T-cell immunotherapy (Peng et al. 2016). In addition, alternative splice variants of *PTEN* have been shown to phenocopy *PTEN* loss (Breuksch et al. 2018). SPLASH2 identified an anchor in PTEN (in 24 cell lines), where one extendor represents novel splicing involving a cryptic exon in the intron between exons 3 and 4; and the other represents annotated splicing between exons 3 and 4 but with two mismatches: an A to G in exon 4 and a T to Aat the boundary of exon 4 (Figure 3B). These two splice junctions are mutually exclusive: cell lines expressing the cryptic exon do not express the annotated junction with two mutations (Figure 3B). This establishes SPLASH2’s distinctive statistical power to jointly detect splicing and multiple genome mismatches, offering potential new markers of *PTEN* loss, and more broadly to identify regulated splicing and sequence variation absent from the reference genome. To the best of our knowledge, this is the first report of such splicing in this critical regulatory gene.

**Figure 3.**
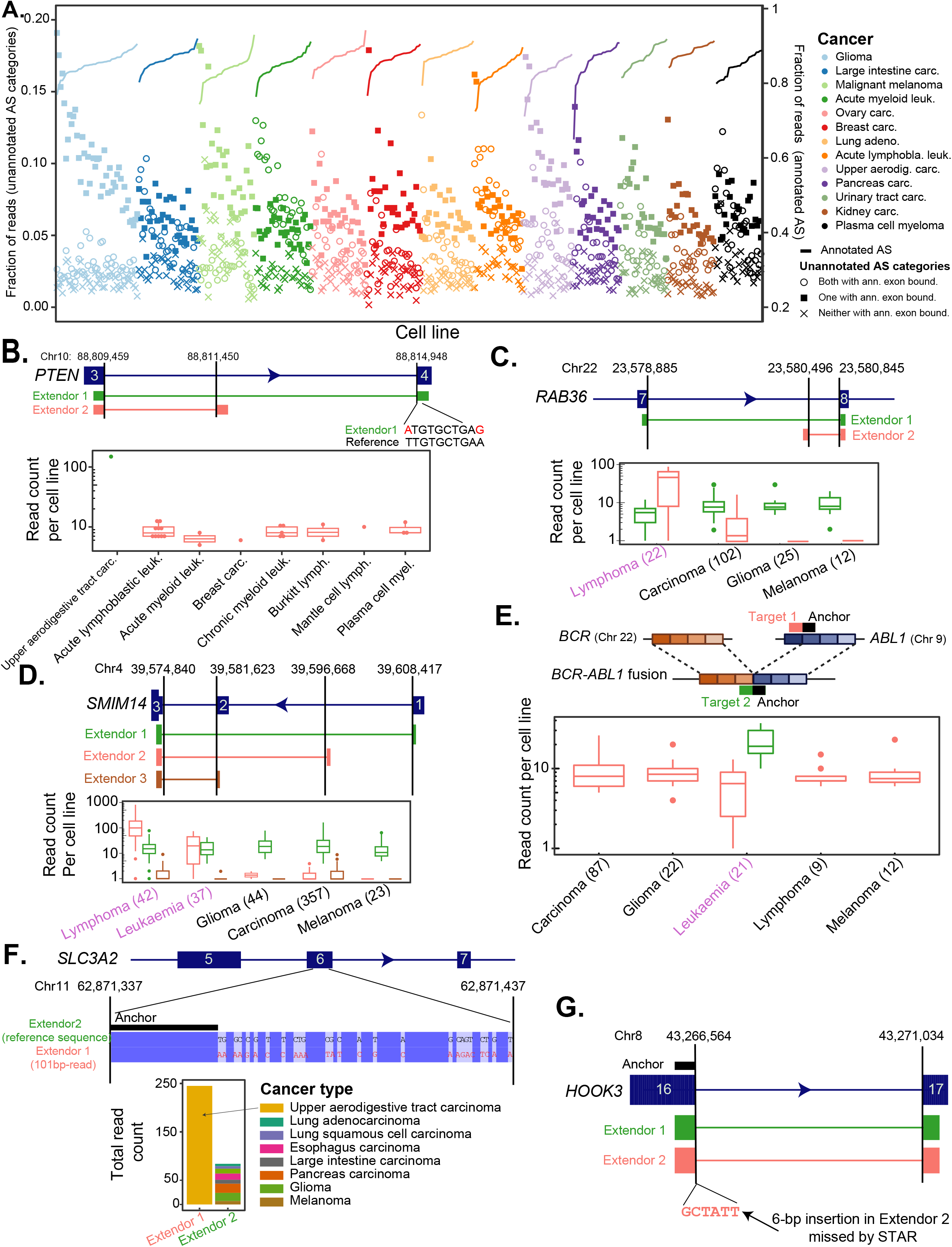
SPLASH2 enables alternative splicing analysis at scale in cancer cell lines. (A) The plot shows the fraction of reads for annotated alternative splicing (lines, right axis) and 3 categories of unannotated alternative splicing (dots, left axis) for each cell line, indicating the majority of detected alternative splicing events are annotated. Cell lines are sorted by cancer and within each cancer by the fraction of reads for annotated alternative splicing. (B) Simultaneous detection of two mutations in an annotated junction (one internal and one at the splice site) and a novel splice junction for *PTEN*. (C) Identification of a cryptic exon for *RAB36* with higher expression than canonical splicing in lymphoma cell lines. Numbers beside each cancer type indicate the number of expressing cell lines. (D) SPLASH2 detected novel cryptic splicing for *SMIM14* with higher expression in lymphoma and leukemia cell lines. (E) Automatic rediscovery of *BCR*-*ABL1* gene fusion using SPLASH2 through an anchor mapping to gene ABL1 with the second extendor indicating the fusion junction, solely expressed in leukemia cell lines. (F) Identification of an anchor for *SLC3A2* exhibiting a high fraction of unaligned reads by STAR, exclusively present in upper aerodigestive tract carcinoma. Multiple sequence alignment of the reference sequence (represented by Extendor 2) and a long 101-bp read for extendor 1 indicates that extendor 1 is due to hypermutation in this gene. (G) Detection of a 6-bp insertion at the splice site of exon 16 in *HOOK3*, reflected in the second extendor of an anchor unaligned by STAR.

We also detected a cancer-type-specific cryptic exon of *RAB36* in lymphoma cell lines (21/22 lymphoma cell lines with expression for the anchor) (Figure 2D). The overexpression of *RAB36*, a member of the RAS oncogene family, has been previously associated with poor prognosis in leukemia patients (Wang et al. 2021). We also detected a splicing specific to leukemia and lymphoma (21 and 15 cell lines, respectively) in *SMIM14*, a small integral membrane protein (Figure 2E). Although the junction found by SPLASH2 is not part of the T2T CAT+Liftoff annotation, it is included in the NCBI RefSeq annotation. Another example of cancer-type-specific regulation found by SPLASH2 is an intron retention in *CLIP3*, with a significantly higher intron retention rate in glioma and melanoma than in other cancer types (Suppl. Figure 6). The downregulation of *CLIP3* has been linked to radiotherapy resistance in glioblastoma stem-like cells (Kang et al. 2021), implying the importance of *CLIP3* variants to the prognosis of glioma patients.

SPLASH2 was also able to rediscover the *BCR*-*ABL1* gene fusion, where the extendor corresponding to the *BCR*-*ABL1* fusion junction was the dominant isoform expressed in leukemia cell lines (Figure 2F). This is consistent with the established recognition of this fusion as a well-known biomarker and treatment target for chronic myelogenous leukemia (CML) patients (Melo and Barnes 2007). This finding highlights the unique ability of SPLASH2 to jointly detect fusions, cryptic splicing, intron retention, and point mutations–all critical transcriptional changes in cancer.

SPLASH2 discoveries extend to other non-canonical transcriptional aberrations missed by STAR. To illustrate this, we focused on anchors effect sizes > 0.2 where one of the top two extendors failed STAR mapping. This analysis unveiled 13,465 anchors with variations in target sequences missed by reference alignment. The expression pattern in *SLC3A2* illustrates the bias of reference alignment (Figure 2G): while the second most abundant extendor mapped to *SLC3A2*, the most abundant extendor accounting for 75% of total anchor reads and observed in an upper aerodigestive tract carcinoma cell line was unmapped. This extendor, reflecting hypermutation in *SLC3A2*, highlights the biases in current alignment-based methods in identifying one of the signatures of cancer genomes. Among other genes with unaligned extendors was *HOOK3*, known to be altered in 0.1% of carcinoma cancers (André et al. 2017). The second extendor of this gene was unaligned by STAR and multiple sequence alignment revealed a 6-bp GCTATT insertion (Figure 2H).

In summary, SPLASH2 is a highly efficient, general analytic framework for biological discovery using genomics sequencing data. In addition to improving the performance of common specialized bioinformatic tools, it discovers novel mutations, transcribed structural variation, and post-transcriptional regulation that may be completely missed by alignment-based methods. To date, the scale of this missing variation, which may be extensive in tumors, has been difficult or impossible to measure in large-scale studies. SPLASH2’s scale enables a next-generation of massive genomic data analysis, including for essentially any RNA or DNA sequencing protocol. Finally, we should point out that SPLASH2 can be extended to long reads (such as PacBio), which will be included in the future release of the software.

## Acknowledgments

We thank Aaron Gitler and Yi Zeng for their suggestions to analyze the ALS data and for useful discussion. J.S. is supported by the National Institute of General Medical Sciences grants R35 GM139517 and the Chan Zuckerberg Data Insights. TZB was funded in part by the Stanford Graduate Fellowship, the NSF GRFP, and the Eric and Wendy Schmidt Center at the Broad Institute of MIT and Harvard.

## Figures

**Suppl. Figure 1.**
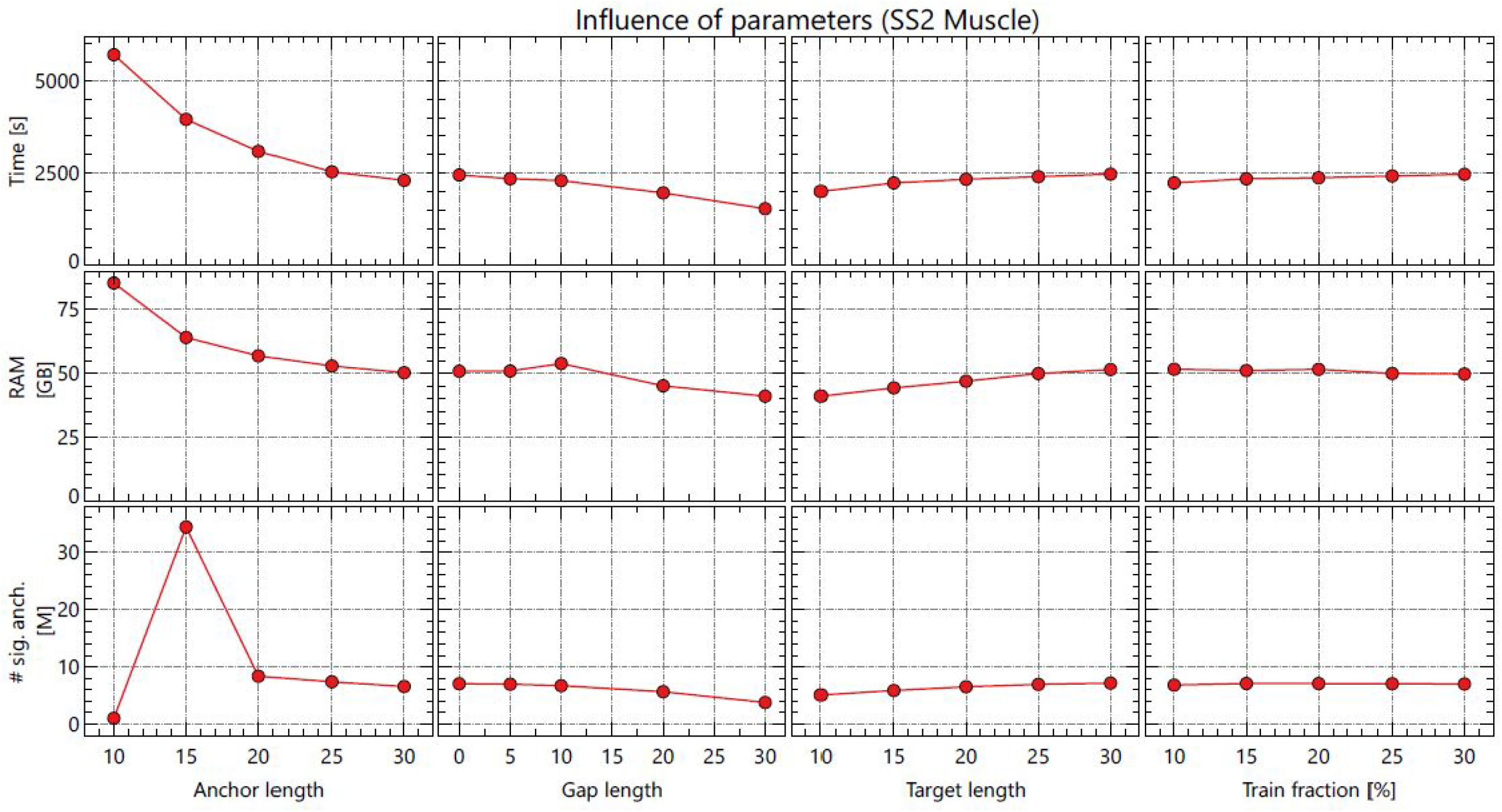
Varying SPLASH2 parameters such as anchor, gap, and target length as well as fraction of training data have negligible impact on run time, RAM usage, and the number of detected significant anchors.

**Suppl. Figure 2.**
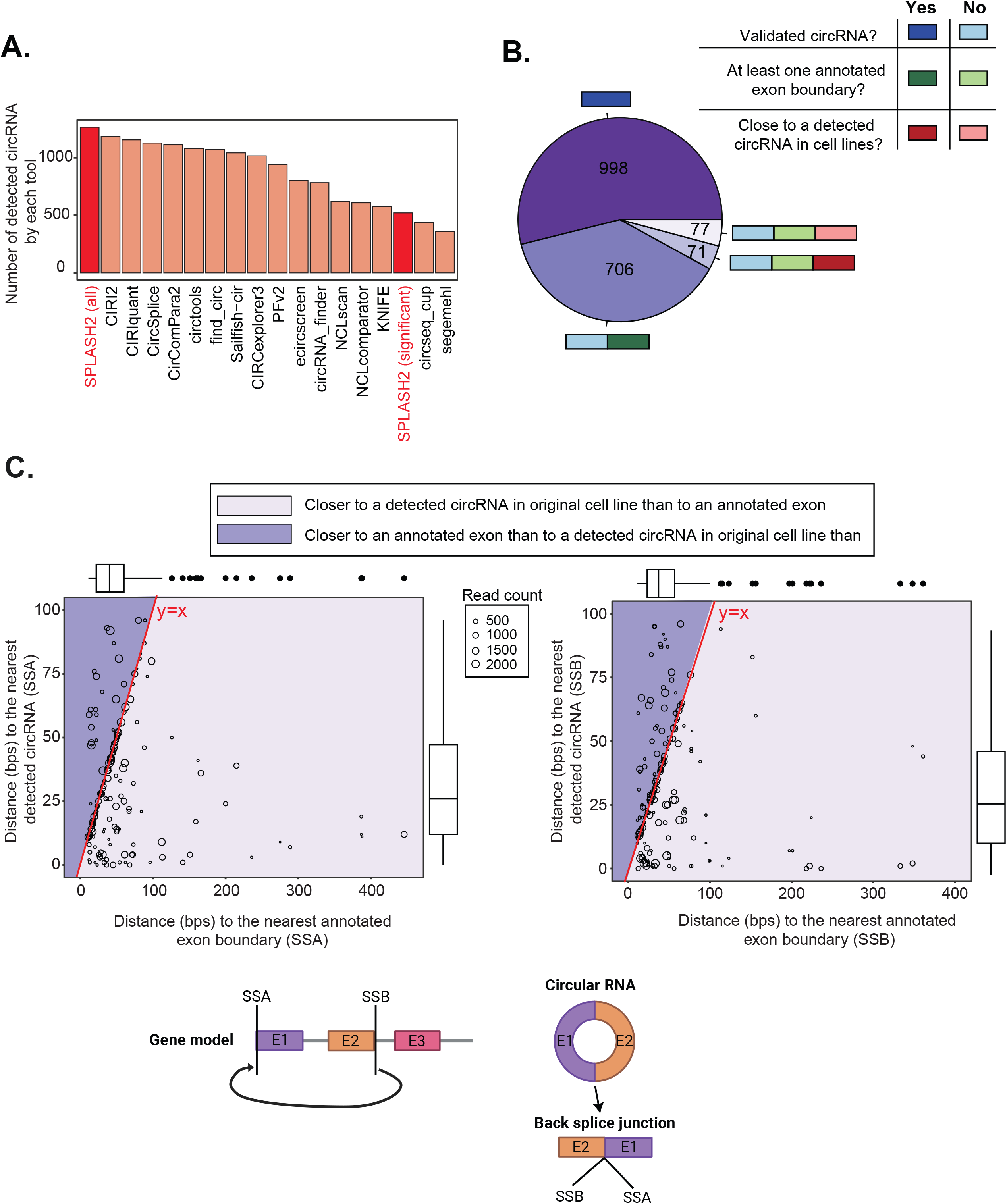
(A) Comparison of the number of validated circular RNAs detected by various tools and SPLASH2, considering all anchors (SPLASH2-all) and only those with significant p-values (SPLASH2-significant). (B) Specificity assessment of SPLASH2’s circular RNA detection on amplicon datasets using validated circRNAs, involvement of annotated exon boundaries, and proximity to detected circRNAs in original cell lines as indicators of true expression. (C) Scatter plots showing the distances of SPLASH2-called circRNAs in amplicon data that are not validated and not at annotated exon boundaries to the nearest annotated exon boundary (x-axis) and the nearest detected circRNA in original cell lines (y-axis) based on the 5’ splice site (SSA) and 3’ splice site (SSB), suggesting the evidence of circRNA expression in original cell lines near these circRNAs.

**Suppl. Figure 3.**
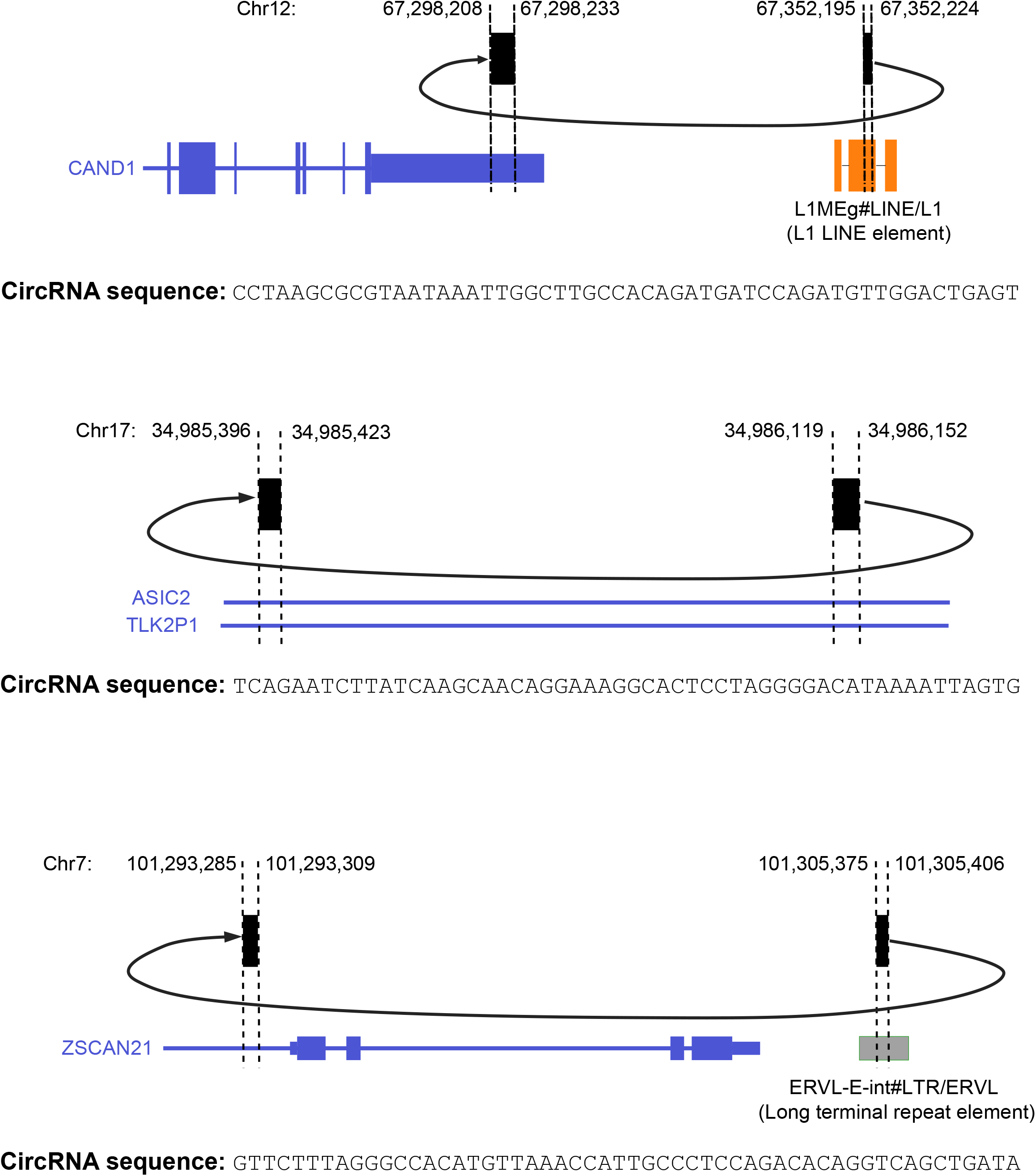
BLAT alignment illustrating the alignment for 3 circRNA sequences called by SPLASH2 as significant in amplicon data, indicating their true mapping to the back splice junctions. These were the only SPLASH2-called circRNAs in amplicon data with a distance of >100 bps from the nearest annotated exon boundaries and also detected circRNAs in the original cell lines detected by other tools.

**Suppl. Figure 4.**
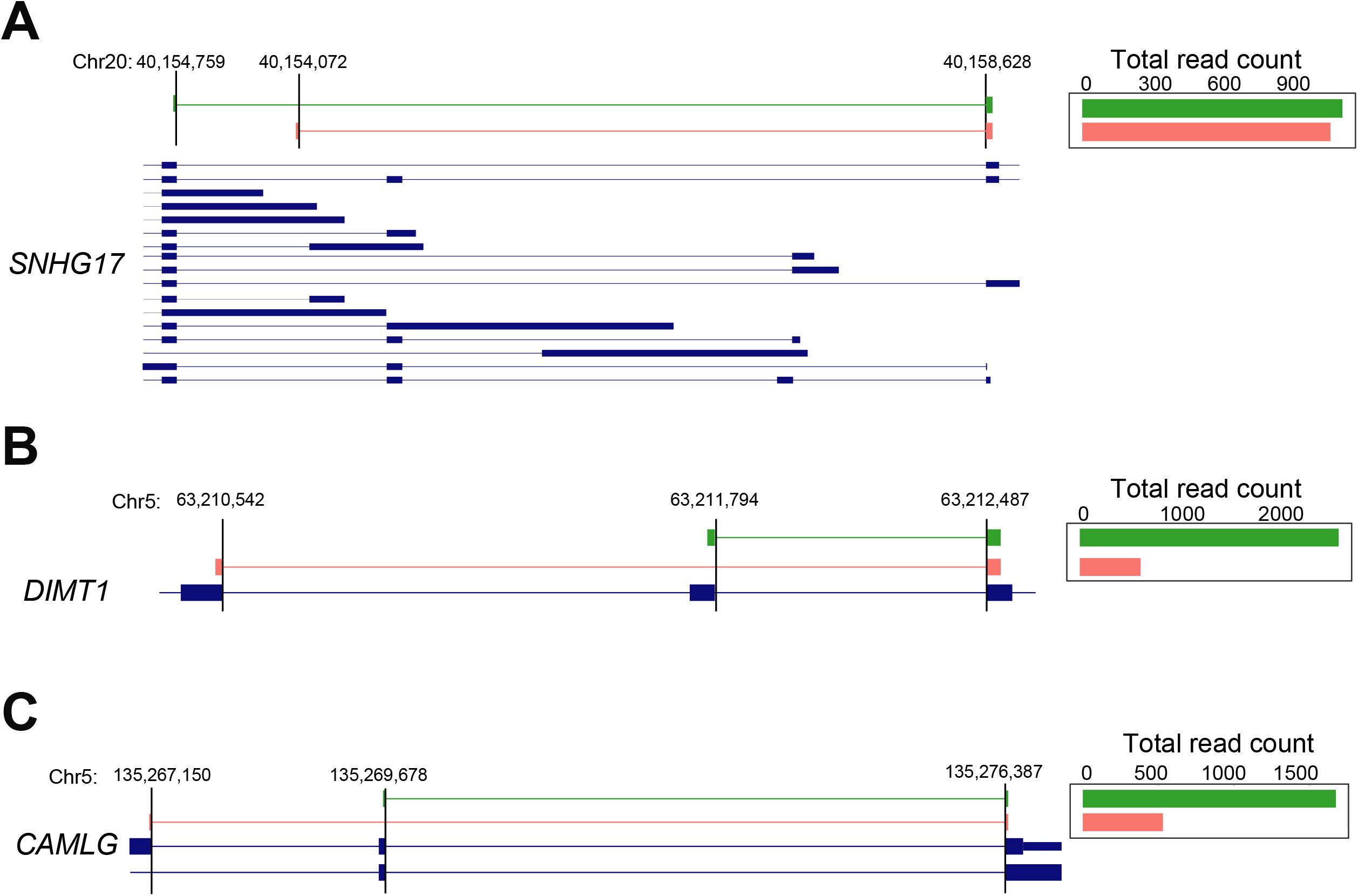
SPLASH2 analysis reveals extensive unannotated alternative splicing in SS2 muscle cells including in genes (A) *SNHG17*, (B) *DIMT1*, and (C) *CAMLG* found in the highest number of cells. Bar plots show the total read count for each annotated (green) and unannotated (red) junction across all expressing cells.

**Suppl. Figure 5.**
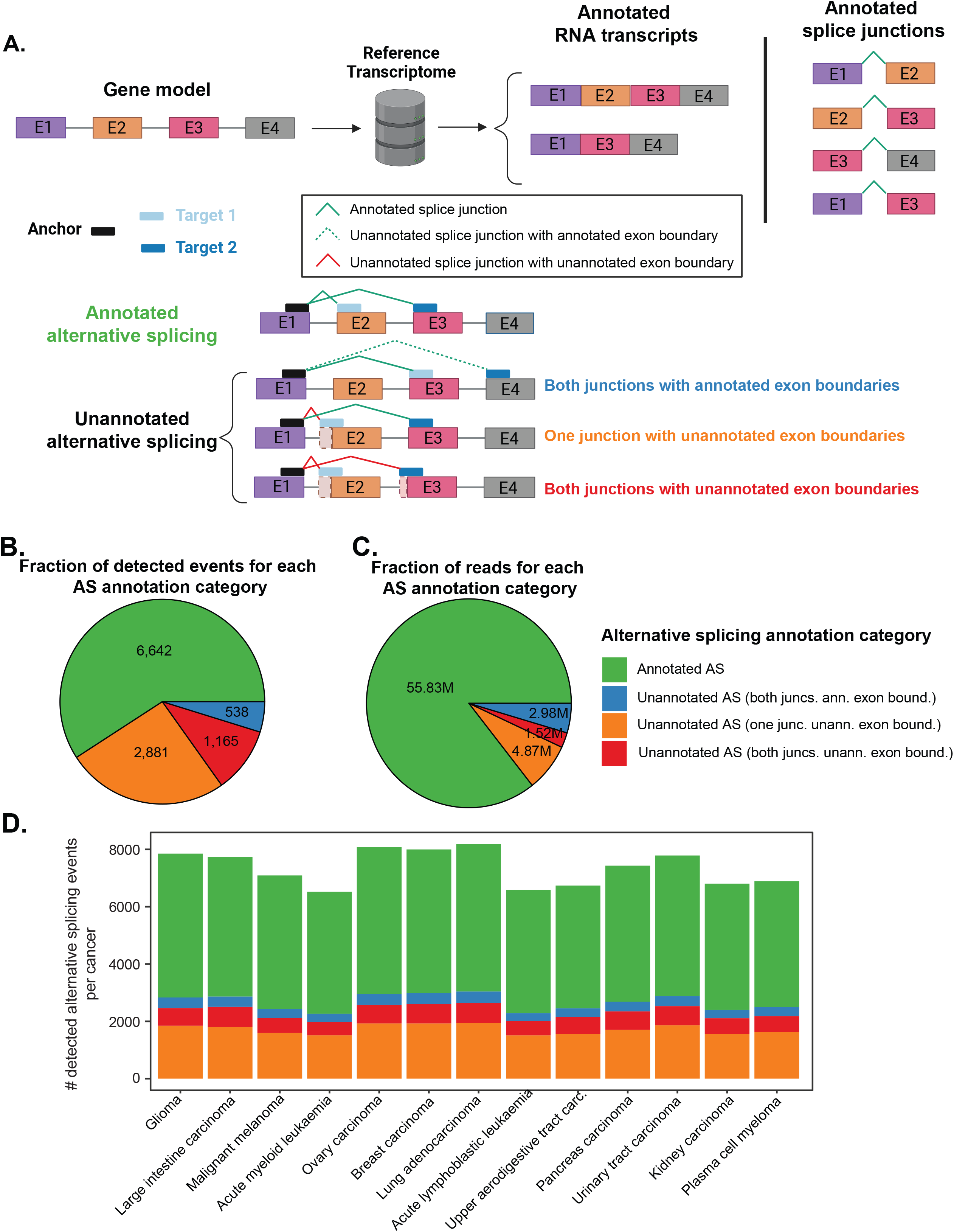
(A) Diagram illustrating the determination of annotation status for detected alternative splicing by SPLASH2 in an example gene with four exons (E1 to E4) and two annotated RNA isoforms (E1-E2-E3-E4 and E1-E3-E4), corresponding to 4 annotated splice junctions. For annotated alternative splicing, both splice junctions (E1-E2 for target 1) and (E1-E3 for target 2) are part of the reference transcriptome. Unannotated alternative splicing is categorized based on the presence or absence of annotated exon boundaries for each junction involved in alternative splicing. (B) Fraction of each annotation category detected across the entire CCLE. (C) Fraction of reads corresponding to each annotation category across the entire CCLE. (D) Number of unique alternative splicing events categorized by annotation status detected by SPLASH2 for each cancer type.

**Suppl. Figure 6.**
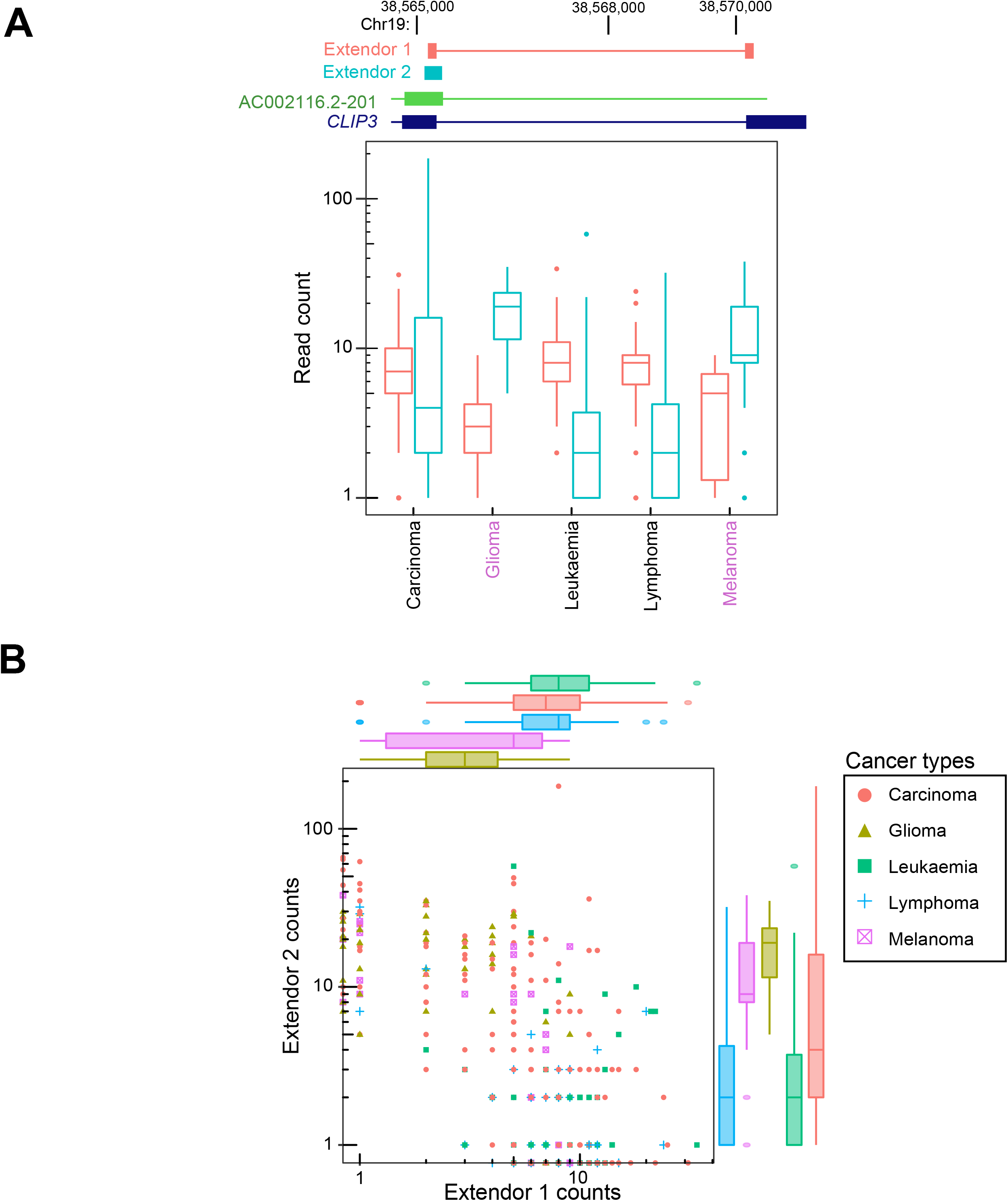
Cancer-type-specific intron retention in *CLIP3*, with a significantly higher intron retention rate in glioma and melanoma than other cancer types. Extendor 1 maps to the splice junction and extendor 2 reflects the intron retention.

## Tables

**Table 1: CircRNA benchmarking**. List of extendors (for both detected and significant anchors) that overlapped with a validated circRNA in circRNA benchmarking study.

**Table 2: RNA mismatches in SS2 muscle cells**. List of anchors with evidence for RNA mismatches called by SPLASH2 in SS2 muscle cells.

**Table 3: CCLE alternative splicing**. List of alternative splicing anchors called by SPLASH2 in the CCLE dataset.

## METHODS

### Code availability

The SPLASH2 pipeline along with detailed instructions and test data are available through a GitHub repository: https://github.com/refresh-bio/SPLASH.

### Data availability

The FASTQ files for the SS2 human muscle cells were downloaded from https://tabula-sapiens-portal.ds.czbiohub.org/. The ALS data was downloaded from the SRA database (SRP185789). The FASTQ files for the 671 cell lines in CCLE from primary tumors were downloaded from the SRA database (SRP186687). The circRNA benchmarking data was downloaded from the SRA database (SRP350843). The FASTA file for human T2T assembly was downloaded from: https://s3-us-west-2.amazonaws.com/human-pangenomics/T2T/CHM13/assemblies/analysis_set/chm13v2.0.fa.gz. Human UCSC GENCODEv35 CAT/Liftoff v2 transcriptome annotation file was downloaded from: https://s3-us-west-2.amazonaws.com/human-pangenomics/T2T/CHM13/assemblies/annotation/chm13.draft_v2.0.gene_annotation.gff3

### Hardware

Run time and memory consumption were evaluated on a machine equipped with AMD 3995WX 64-Cores CPU, 512 GB RAM, and 4 HDDs configured in RAID5.

### Anchor target counting in SPLASH2

One of the most time-consuming parts of the first version of SPLASH was related to the retrieval of anchor-target pairs with counts per sample. While there are efficient *k*-mer counting tools such as Jellyfish (Marçais and Kingsford 2011), KMC (Kokot, Dlugosz, and Deorowicz 2017), and DSK (Rizk, Lavenier, and Chikhi 2013), they all fall short of counting gapped *k*-mers that consist of two segments with gap (relative to the input sequence) between them and thereby cannot be directly adopted for SPLASH, which by design allows non-zero-length gaps between anchors and targets. Given SPLASH requires a contingency table for each unique anchor, collecting the targets across all samples for each anchor efficiently and with minimal memory usage is crucial, posing a notable algorithmic challenge. SPLASH2 resolves this by enumerating all (*a+g+t*)-mers (*a, g, t* being anchor, gap, and target lengths, respectively) with their counts using KMC. Then these sequences are lexicographically sorted with KMC-tools (Deorowicz, Debudaj-Grabysz, and Grabowski 2013; Kokot, Dlugosz, and Deorowicz 2017). This leads to adjacent occurrences of unique anchors, enabling efficient gap removal and the collapsing of unique targets in the subsequent step via a linear traversal over the (*a*+*g*+*t*)-mers. This process collects all targets for the current anchor, with gaps being omitted. Subsequently, targets are sorted, unique targets are merged, and their counts are updated. We note that this scheme heavily utilizes the sorted order of *(a+g+t)*-mers. Using hash table-based *k*-mer counters (like Jellyfish or DSK) would make the processing much harder (if possible).

### SATC file format

The pipeline is implemented as a set of separate modules, communicating through specialized files with the SATC (**S**ample ID, **A**nchor **T**arget **C**ount) format. Each SATC file stores a list of records consisting of sample ID, anchor sequence, target sequence, and count (i.e., each record gives the read count for a pair of anchor-target in the sample specified by sample ID). For each field of a record, a minimal number of bytes is used; however, to minimize the file size some additional techniques are employed. After the first step of SPLASH2, the SATC file contains a single sample ID and is sorted relative to the anchor and then targets within each anchor. This arrangement results in adjacent records sharing a certain number of the most significant bytes. Typically, the bytes used to represent the sample ID are identical, and often the next bytes are also the same. To optimize storage, we prepend an extra byte to each record, indicating the number of identical leading bytes compared to the preceding record. This strategy results in the removal of redundant bytes, enabling the allocation of the minimum number of bytes per record. During the reading, these bytes are reconstructed based on the previous record. We also applied Zstandard (https://github.com/facebook/zstd), a general-purpose compression algorithm, to further compress the SATC files.

### Merging SATC files

Step 1 generates SATC files as many as the number of samples. Depending on the dataset, hundreds of input samples/streams may exist. K-way merging is a known problem in computer science, where the goal is to merge multiple sorted input sequences into a single sorted output sequence, and there are various methods to handle it. In SPLASH2, we used the binary heap K-way merging approach. For each unique anchor, we build a min-heap of size equal to the number of input samples for its targets. When the user specifies the maximum number of targets used for the contingency table (defined by --n_most_freq_targets_for_stats parameter), we employ another binary heap where each element is the anchor’s target, total target count across all samples, and the list of sample IDs with counts. This binary heap is a max-heap with respect to the total target count of size equal to the specified number of most frequent targets to be kept. This approach reduces RAM requirements by discarding less frequent targets. After merging records for each anchor, the contingency table is constructed, and all statistical computations are performed. This approach enhances runtime efficiency and reduces disk space requirements by eliminating the need to store merged data for each anchor, as only the output statistics are saved in the output file.

### The architecture of the SPLASH2 code

SPLASH2 comprises a suite of applications coded in the C++ programming language, accompanied by a single Python script serving as a wrapper to manage their execution. In the first step, we leverage KMC and KMC-tools, both being parallelized. Parallelization of other steps is implemented at a higher level where the Python wrapper script creates separate processes to run multiple instances of each remaining single-threaded executable in parallel. To be able to run multiple instances of SATC merging processes, SPLASH2 distributes records of a single sample into multiple SATC files, referred to as bins, separated based on the hash values of the anchors. The single merging process reads the given bin number of all samples. Three parameters characterize the parallel configuration of SPLASH2: `--n_threads_stage_1`, `--n_threads_stage_1_internal`, and `--n_threads_stage_2.` In the first stage, a maximum of `n_threads_stage_1` processes can be run simultaneously, each internally utilizing up to `n_threads_stage_1_internal` threads for (*a*+*g*+*t*)-mers counting, (*a*+*g*+*t*)-mers sorting, gap removal, and hash-based anchor distribution to bins. In the second stage, a maximum of `--n_threads_stage_2` processes can be run at once, with each process internally using one thread. Merging and statistical computations are performed through a single executable. Therefore, the maximum number of CPU cores simultaneously used by the pipeline is determined by MAX(`n_threads_stage_1 × n_threads_stage_1_internal,n_threads_stage_2`). If *p* cores are available, we recommend the configuration where `n_threads_stage_2` is set to *p* and `n_threads_stage_1` and `n_threads_stage_1_internal` are set such that their multiplication equals *p*. When the number of input samples is small, we recommend allocating fewer cores to `n_threads_stage_1` and more cores to `n_threads_stage_1_internal`.

### SPLASH’s statistical methodology

By default, SPLASH2 computes several measures for the interpretability and processing of anchors, such as the average Hamming distance between target sequences, and target entropy quantifying the probability distribution of the targets. The optimization procedure for the p-value computation identifies latent structure in the matrix by splitting the counts matrix into train and test portions, computing 1-dimensional row and column embeddings (*f* and *c*) respectively (original problem is bi-convex so this is performed via alternating maximization). These *f* and *c* are then used, with the held-out test data, to yield a statistically valid p-value, and compute an effect size measure, which indicates how well the samples can be separated into two groups. Additional outputs such as an asymptotically-valid p-value are also available. Additional statistical details regarding SPLASH are available in (Baharav, Tse, and Salzman 2024). For each anchor, SPLASH computes an “effect size”, which falls within the range of 0 to 1. An effect size of 0 indicates that the target distribution is equivalent between the two input sample groups, while a value of 1 indicates complete disjointedness in the sets of targets across the two sample groups.

### Anchors filtering

To lower the memory requirements and computation time, several filters are applied in SPLASH2 to remove potential artifacts. The first pair of filters are applied in the first step to discard all the anchors for a given sample where: a) its total count is not greater than 5, or b) the anchor contains a poly(A/C/G/T) string of length at least 8. In the second step, all the anchors fulfilling at least one of the following conditions are removed: a) the sum of counts across all samples and targets is not greater than 50, b) the number of unique targets is not greater than 1, or c) the number of unique samples is not greater than 1. All the given values are defaults and can be redefined by the user. In some cases, the number of targets for a single anchor may be large, while only the most abundant ones are relevant. SPLASH2 allows keeping only a user-defined number of most frequent targets per anchor (by default, all targets are used).

### Efficiency evaluation of SPLASH2

We assessed the performance improvement achieved in SPLASH2 relative to the original SPLASH implementation by running both SPLASH2 and SPLASH on two cell lines CCK-81 (SRR8615251) and HS616T (SRR8615424) from the CCLE dataset. We ran the original SPLASH implemented in Nextflow (https://github.com/salzman-lab/SPLASH) using the following command:

~~~
-profile singularity -r main -latest --input input-test.csv
--num_reads_first_pass 2000000000 --num_reads_second_pass 2000000000
--use_read_length false --lookahead 0 --window_slide 1 --run_pvals_only.
~~~

For the SPLASH2, we use default parameters. The run time for SPLASH was 26 hours 38 minutes, while SPLASH2 was completed in 3.5 minutes, suggesting that SPLASH2 was ∼450 times faster than SPLASH1.

For efficiency analysis based on the SS2 muscle cells (1,553 FASTQ files of average size 657MB), KMC was run in RAM-only mode and 16 processes were assigned to the first step (2 threads for each process). For the ALS dataset, KMC was run in the default disk mode and its RAM limit was set to 6GB and 2 processes (each with 16 threads) were assigned to the first step. For the second step, we used 32 computing processes.

Increasing gap length slightly lowers all the measured values, as the number of (*a*+*g*+*t*)-mers decreases. In SS2 muscle cells, the most resource-demanding was the second step (Figure 1B), while in the case of large input samples (i.e., ALS dataset) the first step was more resource-demanding (Figure 1B).

To assess the impact of the input data size (both the number of samples and the size of each sample), we ran SPLASH2 on different numbers of SS2 muscle cells (Figure 1D). We also ran on all ALS samples, subsampling them to specific read counts (Figure 1E). In both cases, run time grows linearly with the input size. In the case of SS2 muscle cells, memory usage also grows linearly, albeit negligible compared to the input size. For ALS, the memory grows linearly with the number of input reads but then saturates at about 300M reads. To evaluate the runtime and memory requirements of STAR, we executed it using 32 threads, analyzing one sample at a time to match the computational resources utilized for SPLASH2 analyses.

### SPLASH runs for SS2 muscle and CCLE analysis

For analysis based on both SS2 muscle cells and CCLE cell lines, we used the following parameters for running SPLASH2:

~~~
--anchor_len 27 --target_len 27 --gap_len 0 --poly_ACGT_len 6
--anchor_count_threshold 50 --anchor_samples_threshold 1
--anchor_sample_counts_threshold 5 --train_fraction 0.2 --fdr_threshold 0.05
~~~

In other words, we set both anchor and target length to 27, and gap length to 0. Anchors containing stretches of A/G/C/T longer than 6 base pairs were excluded. We kept anchors with >50 reads in > 5 input samples (cells). 20% of reads were utilized for the training phase for each anchor. The False Discovery Rate (FDR) threshold for calling significant anchors was set at 0.05.

### CircRNA benchmarking analysis

We ran SPLASH2 on the RNase R +/- samples from each cell line in the circRNA benchmarking study (Vromman et al. 2023). Treatment with RNase R is an experimental procedure to enrich for circRNAs as RNase R selectively degrades linear poly(A)-tailed RNAs. We concentrated our analysis on the 1,560 “validated circRNAs” that were further validated in the circRNA benchmarking study. When we obtained significant anchors called by SPLASH2 for each RNase R +/- sample pair, we extracted the junctional sequence (20 base pairs on each side of the back splice junction) for each validated circRNA using the genomic coordinates reported in Supplementary Table 3 in (Vromman et al. 2023) and the `getSeq` command from `BSgenome.Hsapiens.UCSC.hg38` package in Bioconductor. The junctional sequences were then used to determine the back splice junction breakpoint within each amplicon sequence for validated circRNAs. The breakpoint position was needed to determine whether or not a mapped extendor to an amplicon sequence overlaps the back splice junction. After obtaining the extendor sequences corresponding to each called anchor, we aligned them to the amplicon sequences using Bowtie (Langmead and Salzberg 2012). We defined an extendor as indicative of circRNA expression if it overlapped the breakpoint position and had a Bowtie alignment score <= -6, or if it had a lower Bowtie alignment score but with an overlap of more than 10 base pairs. The number of circRNAs identified by SPLASH2 (521 circRNAs) was obtained by counting unique validated circRNAs that had at least one aligned extendor meeting the above criteria.

### Specificity evaluation of SPLASH2 using circRNA amplicon data

We used amplicon data from the same circRNA benchmarking study (Vromman et al. 2023). This amplicon sequencing dataset was obtained by constructing primers for 1,560 “validated circRNAs” and includes a pair of RNase R +/- samples for each cell line obtained. Similar to our benchmarking analysis on the original cell lines, for amplicon data we applied SPLASH2 to each pair of RNase R +/- samples per cell line. To identify extendors that could be attributed to circRNAs, we aligned the extendors to the human T2T reference genome using STAR in two-pass mode and with chimeric alignment enabled:

~~~
--twopassMode Basic --alignIntronMax 1000000 --limitOutSJcollapsed 3000000
--chimJunctionOverhangMin 10 --chimSegmentReadGapMax 0 --chimOutJunctionFormat
1 --chimSegmentMin 12 --chimScoreJunctionNonGTAG -4 --chimNonchimScoreDropMin
10 --outSAMtype SAM --chimOutType SeparateSAMold --outSAMunmapped None
--clip3pAdapterSeq AAAAAAAAA --outSAMattributes NH HI AS nM NM
~~~

We classified extendors exhibiting a chimeric alignment where both segments align to the same chromosome and strand orientation but in reverse canonical order (consistent with alignment to a back splice junction) as “circRNA extendors”. This approach identified 1,852 unique chimeric alignments or unique circRNAs. Following the same methodology employed in the previous section for analysis of original cell lines, we aligned these circRNA extendors to the amplicon sequences using Bowtie and then followed the same criteria of overlap and alignment score to determine whether the circRNA extendor maps to the back splice junction in the amplicon sequence. Subsequently, 998 circRNA extendors were mapped to validated circRNAs. We then investigated whether the back splice sites for each remaining circRNA extendor (that were not among validated circRNAs) coincided with annotated exon boundaries. Among the remaining circRNA extendors, 706 extendors had at least one back splice site within a distance of <20 base pairs from an annotated exon boundary (Suppl. Fig. 2B). For the remaining 148 circRNA, we compared their coordinates with those of the circRNAs detected by other circRNA tools in the original cell lines as reported in Supplementary Table 2 in (Vromman et al. 2023). Of these, 71 circRNA extendors were found to be close to a detected circRNA, defined as a distance of <20 base pairs (based on either comparing the coordinates of 5’ splice sites or coordinates of 3’ splice sites) (Suppl. Fig. 2B). Finally, among the remaining 77 circRNAs, only 3 circRNA extendors had a distance of >100 base pairs from both closest annotated exon boundaries and detected circRNAs in original cell lines. We blatted (https://genome.ucsc.edu/cgi-bin/hgBlat?command=start) the sequence of these 3 circRNA extendors which showed perfect alignment to back splice junctions (Suppl. Figure 3).

### Splicing anchor analysis

We performed a *post facto* step to identify anchors where the diversity of target sequences could be attributed to alternative splicing. For each anchor, we concatenated its sequence with the sequence of each one of its corresponding targets to form extendor sequences. As we used k=27 for both anchor and target lengths for our analysis, each extendor has 54 base pairs. The extendors were then aligned to the human reference genome (T2T) using STAR. An anchor is categorized as a splicing anchor if STAR detects a splice junction for at least one of its two extendor sequences, corresponding to the two most abundant targets.

For both muscle SS2 and CCLE splicing analysis, we considered those splicing anchors with SPLASH effect_size >0.2 that are in >10 cell lines (for CCLE) or cells (for SS2 muscle).

For a splicing anchor, if STAR reports a splice junction for both of the two extendors, we classify the annotation category for the corresponding alternative splicing by comparing it against the set of annotated splice junctions and exon boundaries in the reference transcriptome (Suppl. Figure 5A). If both splice junctions are annotated in the reference transcriptome, the alternative splicing is categorized as “annotated”. However, if at least one of the splice junctions is not annotated, the alternative splicing is classified as “unannotated” which is further divided into three distinct subcategories based on whether none, one, or both of the splice junctions involve unannotated exon boundaries (Suppl. Figure 5A).

### RNA mismatch classification

If all the base pair substitutions between the targets of an anchor can be performed by a specific mapping, we assign an “RNA mismatch” category to the anchor (Figure 2A). For example, we assign a mismatch category “A<->G” to an anchor, if converting all G’s in its target sequences to A results in a unique sequence. We assign 6 different categories: A<->G, A<->T, A<->C, G<->C, C<->T, and T<->G. We should note when sequencing data is not strand-specific, some of these categories are equivalent and can be merged, such as merging A<->G with C<->T and A<->C with T<->G. Given that the SmartSeq2 protocol is not strand-specific, we report 4 mismatch categories for SS2 muscle cells.

We used the following model to calculate the probability of observing 4 targets with >5% abundance under a biallelic model. As in a diploid genome two alleles are present, observing non-modified or edited RNA where the anchor is adjacent to a bi-allelic SNP should result in observing two targets with high abundance. To compute the probability of observing 4 targets, each with >5% abundance, we bound this probability by considering the probability that either allele has a second target with >5% abundance. We assume that there is only 1 allele and compute the probability of observing a target with > 5% abundance. There are 27*3 different 1-base pair changes from the dominant target, which we assume to be an un-modified allele. The first-order approximation of the probability of an observed target differing from the dominant allele is that the target has one mismatch and if the sequencing error is 1%, the probability of observing this target is less than 0.01/3. We then compute, with n.trials=27*3 to estimate the probability of observing one target with > 5% abundance among all anchor counts as follows:

~~~
sum(rbinom(n.trials,N,(.01/3))/N>.05)/n.trials,
~~~

where N is the number of observations per anchor. Setting N to a range from 100 to 1000 yields a value of 0 (up to computational precision).

### Enrichment analysis of COSMIC genes

To compute the p-value for the enrichment of COSMIC genes among genes with unannotated alternative splicing in CCLE, we first counted the number of unique gene names with SPLASH2-called anchors in CCLE (40,133 genes). Using this count as the background, the null probability of a gene with SPLASH2-called anchor being a COSMIC gene was 736/40,133 = 0.0183. Based on this null probability, the exact binomial p-value for the enrichment of COSMIC genes in genes with unannotated alternative splicing was found to be < 2.2e-16. However, for a more conservative approach, we decided to consider 30,000 as the number of background genes, reflecting the number of protein-coding genes in the human reference genome. This adjustment yielded a null probability of 0.024 and an exact binomial p-value of 1.92e-6.

